# Ploidy dynamics in aphid host cells harboring bacterial symbionts

**DOI:** 10.1101/2021.12.03.471054

**Authors:** Tomonari Nozaki, Shuji Shigenobu

## Abstract

Aphids have evolved bacteriocytes or symbiotic host cells that harbor the obligate mutualistic bacterium *Buchnera aphidicola*. Because of the large cell size (approximately 100 μm in diameter) of bacteriocytes and their pivotal role in nutritional symbiosis, researchers have considered that these cells are highly polyploid and assumed that bacteriocyte polyploidy may be essential for the symbiotic relationship between the aphid and the bacterium. However, little is known about the ploidy levels and dynamics of aphid bacteriocytes. Here, we quantitatively analyzed the ploidy levels in the bacteriocytes of the pea-aphid *Acyrthosiphon pisum*. Image-based fluorometry revealed the hyper polyploidy of the bacteriocytes ranging from 16- to 256-ploidy throughout the lifecycle. Bacteriocytes of adult parthenogenetic viviparous females were mainly 64-128C DNA levels, while those of sexual morphs (oviparous females and males) were consisted of 64C, and 32-64C cells, respectively. During post-embryonic development of viviparous females, the ploidy level of bacteriocytes increased substantially, from 16-32C at birth to 128-256C in actively reproducing adults. These results suggest that the ploidy levels are dynamically regulated among phenotypes and during development. Our comprehensive and quantitative data provides a foundation for future studies to understand the functional roles and biological significance of the polyploidy of insect bacteriocytes.

## Introduction

Endopolyploidy or somatic polyploidy is generated through endoreduplication cycles, in which the nuclear genome is repeatedly replicated without mitotic cell division^[1–4]^. Polyploid cells are typically observed in tissues with high metabolic demand and are considered not only to play essential roles in normal development but also to influence ecologically important traits, such as body or organ size, growth rate, and nutrient storage^[5, 6]^. Polyploidization is often attributed to increasing cellular size, metabolic rate, and gene expression levels owing to the increasing availability of DNA templates for transcription; these ideas have been supported by experiments in model organisms^[3, 7, 8]^. Nevertheless, relatively few empirical studies have focused on endopolyploidy in evolutionary and ecological contexts^[6]^.

Many insects, because of their unbalanced diet, live in symbiotic associations with microorganisms, which can provide host insects with nutrients that cannot be synthesized by insects or obtained from their diets^[9–11]^. Such endosymbiotic microorganisms are frequently harbored in insect-specific cells or tissues (endosymbiosis; ^[9, 12–14]^). So far, many researchers have pointed out that symbiotic host cells (mycetocytes or bacteriocytes) have large nuclei and thus, would be polyploid in various insect species^[11–14]^. Considering the well-known effects of polyploidy such as cell enlargement and upregulation of gene expression, we can predict that polyploidization of symbiotic host cells may have critical roles in endosymbiosis between insects and microorganisms; however, very few studies have quantified the ploidy levels of symbiotic host cells in insects. Exceptionally, Nakabachi *et al*. (2010) revealed that in psyllid insects, *Pachypsylla venusta* (Osten-Sacken), symbiotic host cells were 16-ploid in both adult males and females^[15]^. There are other reports on the ploidy levels of symbiotic host cells, but they are incomplete^[16–18]^.

Aphids (Hemiptera: Aphididae) have evolved symbiotic relationships with the gamma-proteobacterium *Buchnera aphidicola*, which supplies the host with vitamins and essential amino acids that aphids cannot synthesize or derive insufficiently from their diet, the plant phloem sap^[10, 19, 20]^. *B. aphidicola* lives within large aphid cells, called “bacteriocytes.” The cells and another type of cells (small and flattened cells, called “sheath cells”) are grouped into organ-like structures, referred to as “bacteriomes” (Figure 1a, b, ^[21]^). Bacteriocytes were long thought to be polyploid because of their large cellular and nuclear sizes (ca. 100 and 20 μm in diameter, respectively)^[12, 13, 22]^. Moreover, aphid bacteriocytes should be highly metabolically active, because they provide *B. aphidicola* with a large amount of essential amino acids and other metabolites that the symbiont can no longer produce, owing to the massive gene losses^[23–25]^. Therefore, polyploidy may be a key element in the maintenance of the symbiotic system, as has been shown in between leguminous plants and rhizobium bacteria^[26–28]^; however, information about the ploidy levels of aphid bacteriocytes remains scarce.

**Figure 1.**
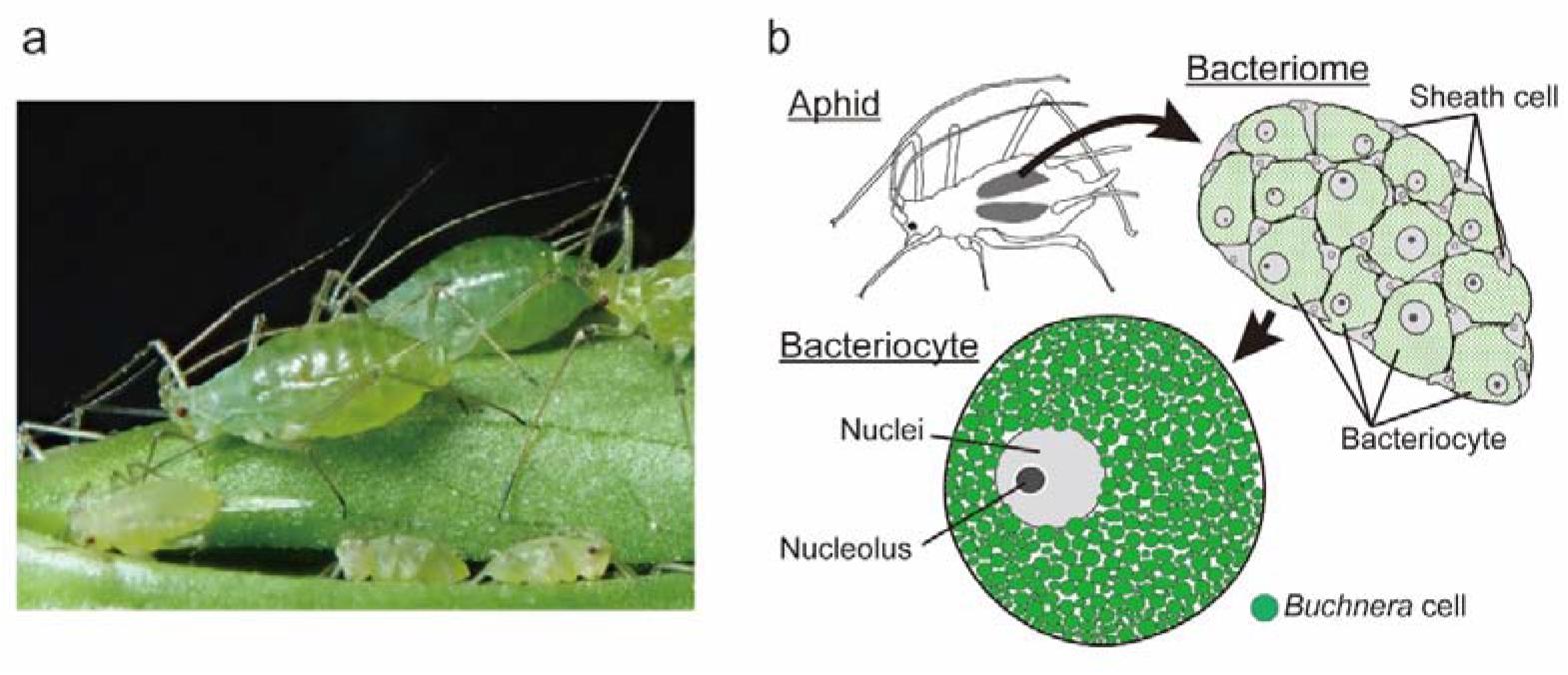
Intracellular symbiosis in the pea-aphid, *Acyrthosiphon pisum*. **a** a photograph of pea-aphids (viviparous females). **b** Aphids harbor their bacterial symbionts in the specialized organ, bacteriome. The organ contains two types of cells, bacteriocytes and sheath cells. Bacteriocytes are symbiotic host cells that harbor *Buchnera aphidicola* in the cytoplasm and are remarkably large (approximately 100 μm in diameter); sheath cells are much smaller than bacteriocytes and do not present *Buchnera*. The function of sheath cells has not been well studied. Illustrations were drawn according to Koga *et al*.^[21]^.

Pea-aphid, *Acyrthosiphon pisum* (Harris), is one of the best-studied aphid species in terms of life history (Figure 2, ^[29, 30]^), polymorphism (Figure S1, ^[30–32]^), embryonic development^[33–35]^, nutritional symbiosis^[23–25, 36–38]^, and genomics^[39–41]^. Detailed descriptions have focused on bacteriocyte development during aphid embryogenesis^[21, 22]^, and it has been revealed that the number and volume of pea-aphid bacteriocytes increase during post-embryonic development (^[38, 42]^ but see ^[43, 44]^). These observations imply that aphid bacteriocytes are already polyploid at the end of embryogenesis and become hyper polyploid during post-embryonic development. However, no quantitative data in this aspect have been reported. It should also be noted that most of the previous studies on aphid bacteriocytes characterized only viviparous insects, excluding other morphs (e.g., oviparous females and males)—presumably because of the ease of rearing viviparous insects in the laboratory (Figure S1).

**Figure 2.**
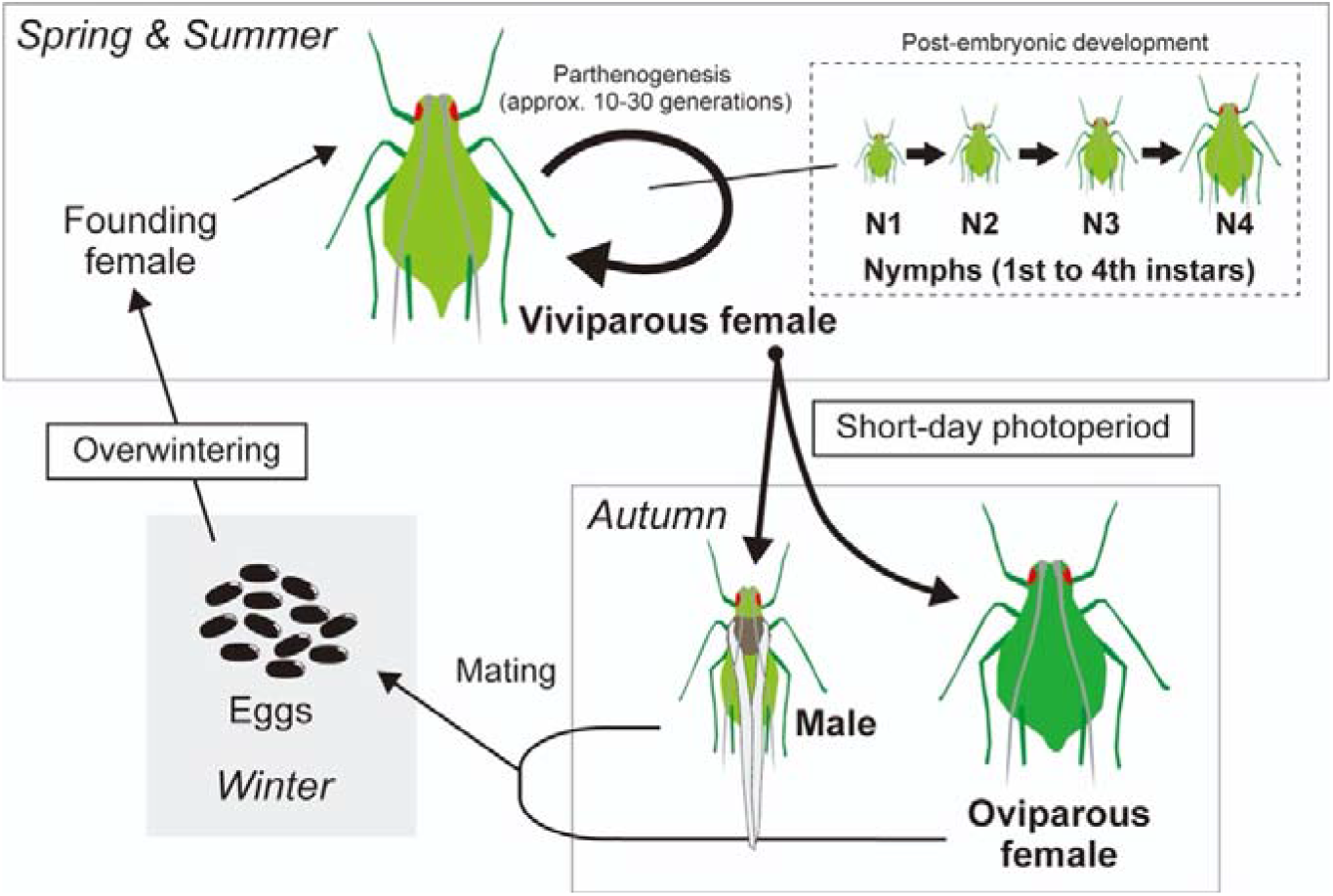
The life cycle and polymorphism of pea-aphids. In the field, viviparous females are observed from spring to summer on their host plants and rapidly reproduce through viviparous parthenogenesis. N1-4 represents the first to fourth instar nymphs. Larviposited nymphs grow up to adults after molting four times. Oviparous females and males emerge in autumn; they are induced by short-day photoperiod. Oviparous females mate with males and produce eggs (sexual reproduction). All aphid morphs harbor *B. aphidicola* inside of their bacteriocytes. Aphid morphs and developmental stages that we have used for ploidy analysis of their bacteriocytes were bold lettered. Illustrations were drawn according to Ogawa & Miura^[30]^.

In this study, we investigated the pattern of polyploidization and the cellular features of pea-aphid bacteriocytes. We first observed the cytological features, such as cell size and nucleolar number/size of bacteriocytes from adults of viviparous/oviparous females and males, and other developmental stages of viviparous insects. Then, using image-based fluorometry established in the present study, we determined the ploidy levels of bacteriocytes and compared these levels among morphs (adult viviparous/oviparous females and males). We also performed ploidy analysis on the cells at each stage of post-embryonic development (from first-instar nymphs to senescent adults in viviparous insects). Finally, based on our observation of the cytological features of bacteriocytes, we discussed the potential effects of bacteriocyte polyploidy on the aphid/*Buchnera* intracellular symbiosis.

## Results

### General observation and methods for ploidy analysis on aphid bacteriome cells

Consistent with previous observations^[9, 21, 22, 40]^, the bacteriome of viviparous aphids consisted of two types of cells: bacteriocytes and sheath cells (Figure 3). Bacteriocytes contained *Buchnera* cells and were much larger than sheath cells. Sheath cells exhibited a flattened morphology and surrounded the bacteriocytes. Both cell types possessed a single nucleus. Bacteriocytes had a single prominent nucleolus, which was not stained using DAPI, but using “Nucleolus Bright Red” staining (Figure 3). Most sheath cells also had a single nucleolus, yet a small number had two. “Nucleolus Bright Red” also stained the peripheral region of *Buchnera*, probably because of the richness of RNA around *Buchnera* cells.

**Figure 3.**
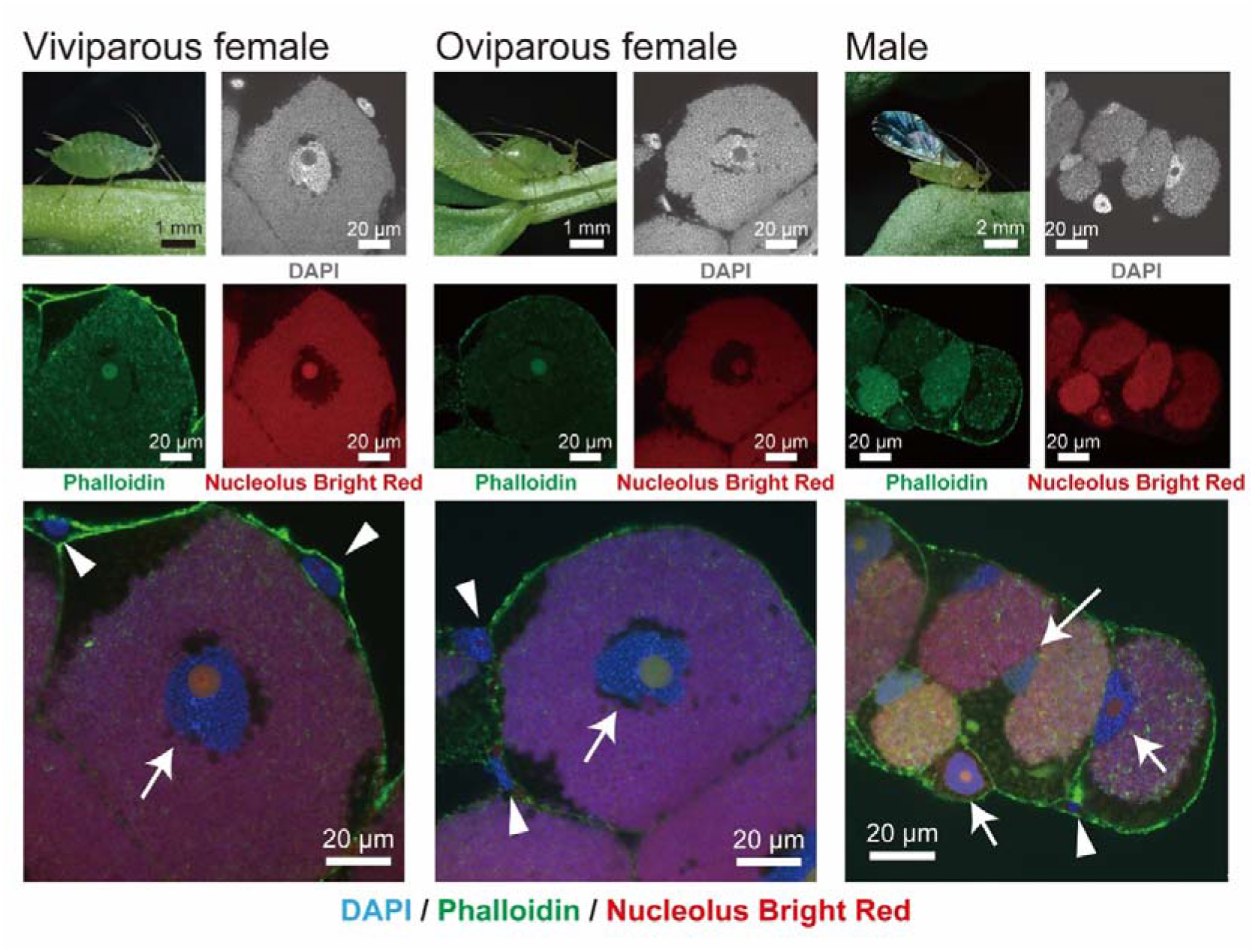
Morphology of bacteriocytes and sheath cells from each morph of aphids visualized using DAPI/Phalloidin/Nucleolus Bright Red staining. DNA and F-actin were stained by DAPI (gray or blue) and Phalloidin (green), respectively. The nucleolus, which is the site of ribosome biogenesis, was visualized by Nucleolus Bright Red (red). This dye binds RNA electrostatically, therefore the cytoplasm of bacteriocytes and *Buchnera* cells were also stained. Bacteriocytes (*white arrows*) had single prominent nucleolus, and the cell sizes were much larger than sheath cells (*white arrowheads*) in all aphid morphs.

To determine the most suitable methods for ploidy analysis of aphid bacteriocytes, three types of methods, flow cytometry, Feulgen densitometry, and fluorometry were compared. First, flow cytometry successfully detected the nuclei of bacteriome cells and heads, and distinct peaks were present (Figure S2). There were several peaks, which can be categorized as ploidy classes based on head peaks, assuming that the smallest peaks correspond to a diploid population. We recognized peaks up to 256C (256-ploidy) cells but could not distinguish cell types (i.e., bacteriocytes or sheath cells) in this method due to a lack of cytological information. Note that “C” means haploid genome size, for example, 2C = diploid and 8C = octoploid. Second, Feulgen densitometry also showed several ploidy levels of up to 128C (Figure S3) in bacteriocytes. Sheath cells mainly consisted of 16-32C cells. However, we found that many cells were lost during the experimental procedures because the number of observed nuclei was too small.

We found the third method, image-based fluorometry for isolated nuclei, the best for quantitative ploidy analysis of aphid bacteriocytes (Figure 4). Fluorometry showed distinct peaks of integrated fluorescence intensity, and they could be categorized as each ploidy class based on the intensity of the smallest peak in head cells (diploid population). The results were consistent with other methods; ploidy levels were 32C-256C in bacteriocytes and 16C-32C in sheath cells. In this analysis, the nucleolus size was used to discriminate between cell types. During cytological observation, we obtained the size distribution of the nucleolus, and it was revealed that the nucleolus of bacteriocytes was always larger than that of sheath cells (Figure S4). Based on the results, we determined the threshold of the size of the nucleolus. More specifically, in viviparous females, nuclei that have nucleoli larger than 20 μm^2^ were categorized into bacteriocytes. Note that the peaks of sheath cells were not distinct or reliable for categorizing their ploidy class; therefore, we showed results focusing on bacteriocytes in the following sections.

**Figure 4.**
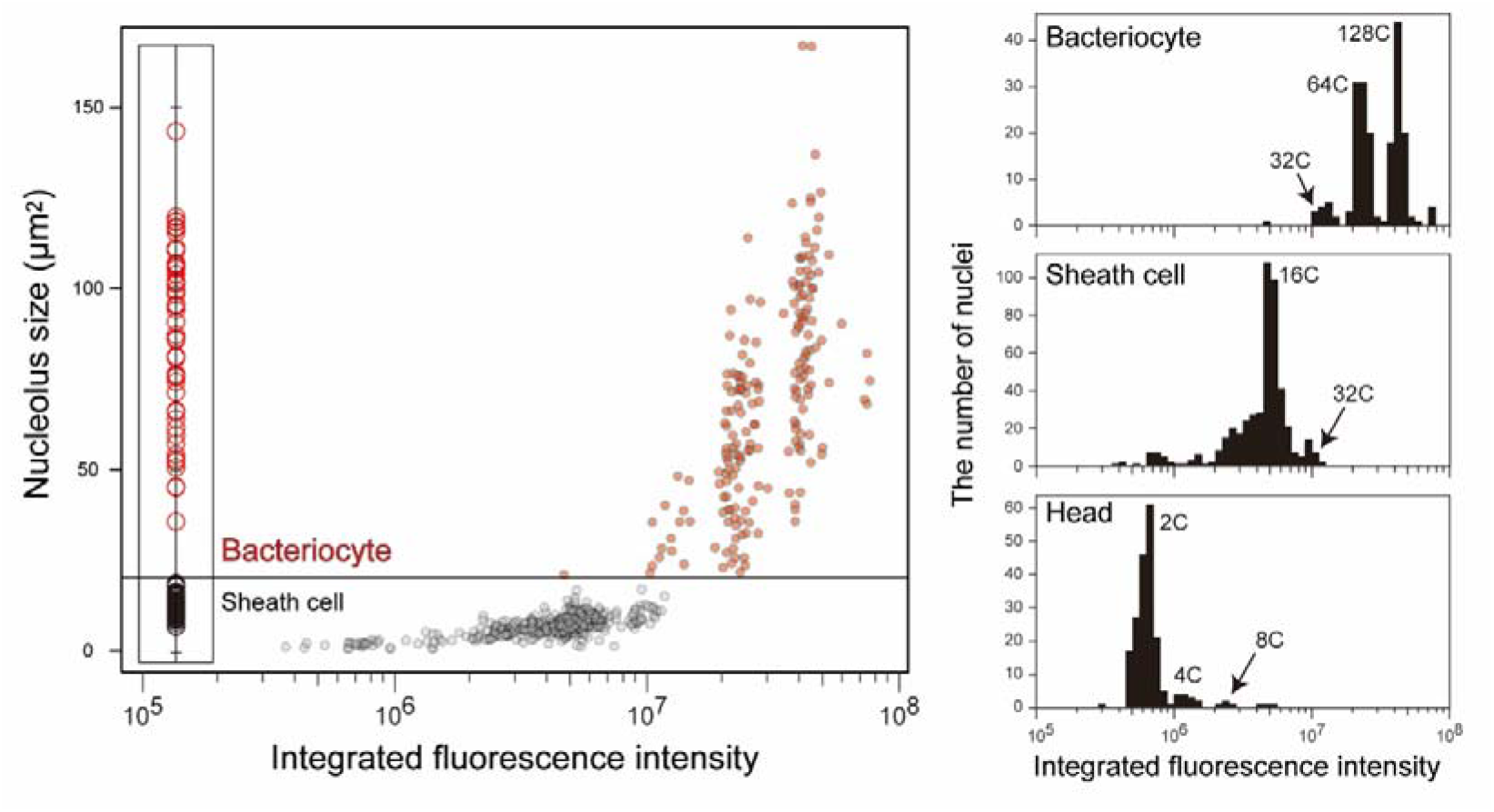
Ploidy analysis of aphid bacteriocytes using DAPI-fluorometry. A representative result from the analysis of adult viviparous females is presented. Isolated nuclei of bacteriome cells were stained using DAPI, image-captured with a CCD camera, and their integrated fluorescence intensity was measured using ImageJ software. Nuclei were categorized into “bacteriocytes” or “sheath cells,” based on the size distribution of nucleolus (see Material and methods). Relative ploidy levels were calculated based on the data from head cells which are mainly diploid. Bacteriocytes of adult viviparous aphids consisted of 16C-256C cells, and 64-128 cells were dominant, while sheath cells exhibited lower ploidy levels (mainly 16C). “C” means haploid genome size, for example, 2C = diploid and 8C = octoploid.

### Cellular features of bacteriome cells in viviparous and oviparous females, and males

The cellular features were generally consistent among adults of three morphs, viviparous and oviparous females, and males (Figure 3). *Buchnera*-absence zones in the cytoplasm of bacteriocytes, which are considered to be degeneration of *Buchnera*^[45]^, and bacteriocytes degeneration^[46]^ were both observed more frequently in male bacteriocytes than in females (Figure 3). The cell size of bacteriocytes was significantly different among morphs (LM with type II test, *F* = 286.15, *df* = 2, *p* < 0.001, Figure S5). Viviparous females had significantly larger bacteriocytes [449847.05 ± 21583.0 (mean ± SEM) μm^3^, *n* = 22] than other types of aphids (oviparous females, 298989.9 ± 16196.6 μm^3^, *n* = 24, and males, 29020.35 ± 3001.6 μm^3^, *n* = 37) (Tukey’s test, *p* < 0.05, Figure S5). The size of nucleoli was significantly different between bacteriocytes and sheath cells, regardless of aphid morphs (LMM with type II test; viviparous females, χ^2^ = 618.4, *df* = 1, *p* < 0.001, oviparous females, χ^2^ = 1,430.4, *df* = 1, *p* < 0.001, males, χ^2^ = 261.37, *df* = 1, *p* < 0.001, Figure S3). There was no overlap in the nucleolus size between cell types (Figure S4). Based on these data, we determined the threshold of the size of the nucleolus to discriminate between bacteriocytes and sheath cells. Specifically, in viviparous and oviparous females, and males, nuclei that have nucleoli larger than 20 μm^2^, 20 μm^2^, 8 μm^2^, were categorized into bacteriocytes, respectively.

### Ploidy analysis on the bacteriocyte of viviparous and oviparous females, and males

Ploidy analysis of the adult bacteriocytes revealed that the cells were highly polyploid (from 32C to 256C) in all phenotypes (Figure 5). We found variation in the level of ploidy; bacteriocytes of viviparous females, oviparous females, and males mainly consisted of 64-128C (45% for each), 64C (70%), and 32-64C (30% and 47%), respectively. There were significant differences in the degree of polyploidy (median ploidy level) in bacteriocytes among the three aphid phenotypes (Brunner–Munzel test with Bonferroni adjustment; viviparous females vs. oviparous females, *p* < 0.05; viviparous females vs. males, *p* < 0.05; oviparous females vs. males, *p* < 0.05; Figure 5).

**Figure 5.**
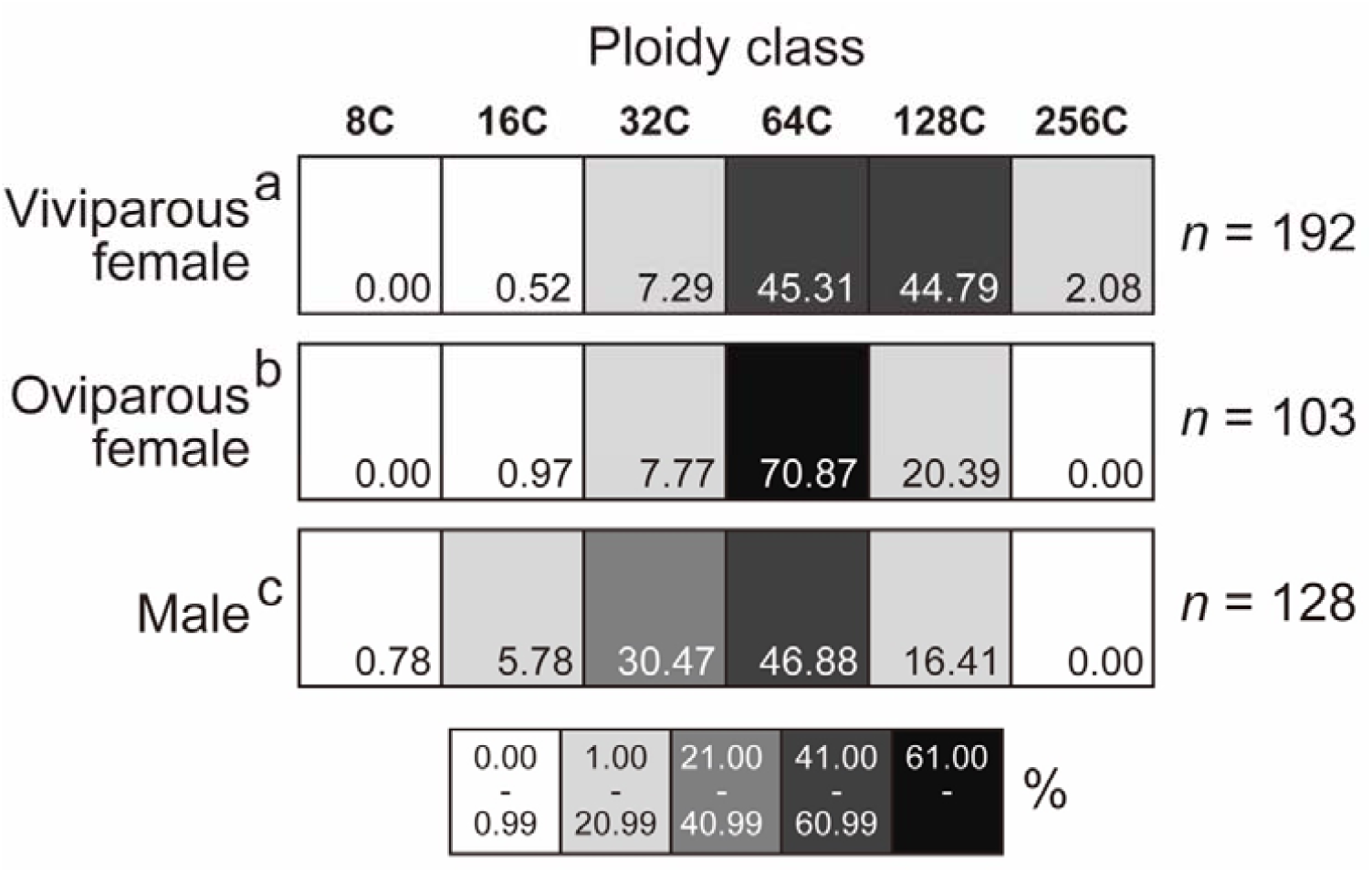
Ploidy distribution of bacteriocytes among aphid morphs, viviparous females, oviparous females, and males. Sample size (numbers of bacteriocyte nuclei) is shown at the right. Different letters with aphid categories indicate significant differences in the median ploidy class (Brunner–Munzel test with Bonferroni adjustment, *p* < 0.05). Bacteriocytes of adult viviparous females in the pea-aphid were highly polyploid and were mainly 64-128C cells. The cells of oviparous females and males were mainly 64C and 32-64C, respectively. The degree of polyploidy levels was highest in viviparous females.

### Fecundity and longevity of viviparous and oviparous females

Viviparous females in this strain laid 95.25 ± 24.75 (mean ± SD, *n* = 12) nymphs during their lifetime, while oviparous females oviposited 28.83 ± 6.52 (*n* = 12) eggs (Figure S6). The total number of nymphs laid by viviparous females was significantly higher than that of eggs oviposited by oviparous females (GLMM with type II test, χ^2^ = 449.74, *df* = 1, *p* < 0.001). Viviparous females lived longer [44.33 ± 4.38 (mean ± SEM) days] than oviparous ones (26.08 ± 2.31 days), yet there was no interaction between female types and their lifetime (GLMM, with type II test; female type, χ^2^ = 118.13, *df* = 1, *p* < 0.001, lifetime, χ^2^ = 69.32, df = 1, *p* < 0.001, and the interaction χ^2^ = 0.74, *df* = 1, *p* = 0.39). Viviparous females started reproducing from days 2-3 and the rate of larviposition peaked during days 3-20 but slowed down during days 21-28. They lived at most 50-55 days, although most of them stopped the larviposition after day 30. In oviparous females, first oviposition and mating with males were observed on days 3-4. They actively laid eggs until day 14, but their death was observed almost simultaneously (Figure. S6).

### Cellular features of bacteriome cells in each stage of post-embryonic development

At 16 °C, viviparous (and apterous) aphids reached adult stage approximately 14 days after birth [13.73 ± 0.32 (mean ± SEM), *n* = 16, Figure 6a]. In particular, N1, N2, N3, and N4 periods lasted for 3.2 ± 0.11, 3.0 ± 0.09, 3.3 ± 0.12, and 4.2 ± 0.14 days (mean ± SEM, *n* = 16, Figure 6a), respectively. Adult aphids started reproducing 2 or 3 days after eclosion (molt for an adult) and continued larviposition for approximately 4 weeks (Figure S6). Based on these data, A7 aphids (7 days after eclosion) could be categorized as actively reproducing individuals. A21 aphids (21 days after eclosion) were categorized as senescent individuals, although they continuously produced offspring. During the nymphal stages of viviparous aphids, the morphology of bacteriome cells was generally consistent; all bacteriocytes and most sheath cells were uninuclear (Figure 6b), but very few of the latter cells had several small nuclei. Notably, there were drastic morphological changes in adult stages; bacteriocyte and sheath cell nuclei of A21 individuals were irregularly shaped in comparison with those of young (A0) and reproducing (A7) individuals. Furthermore, in A21, we frequently observed bacteriocytes in which the signals of DAPI and nucleolus bright red signals on *Buchnera* were weak (Figure 6b). These changes were consistent with symptoms of *Buchnera* degeneration and cell senescence, which have been previously reported ^[45, 46]^. Developmental stages had a significant effect on bacteriocyte size (LMM with type II test, χ^2^ = 338.73, *df* = 6, *p* < 0.001). During post-embryonic development, the size of bacteriocytes consistently increased (Tukey’s test: N1 = N2 = N3 < N4 < A0 = A7 < A21, *p* < 0.05, Figure S7). The size of nucleoli was significantly different between bacteriocytes and sheath cells, regardless of the post-embryonic developmental stages (LMM with type II test; N1, χ^2^ = 891.82, *df* = 1, *p* < 0.001, N2, χ^2^ = 294.04, *df* = 1, *p* < 0.001, N3, χ^2^ = 842.31, *df* = 1, *p* < 0.001, N4, χ^2^ = 817.18, *df* = 1, *p* < 0.001, old adults, χ^2^ = 1,405.6, *df* = 1, *p* < 0.001, Figure S8). There was no overlap in the nucleolus size between cell types (Figure S8). Based on these data and the data from young adults, we determined the threshold of the size of the nucleolus for ploidy analysis (in N1 and N2, 10 μm^2^, and later stages, 25 μm^2^).

**Figure 6.**
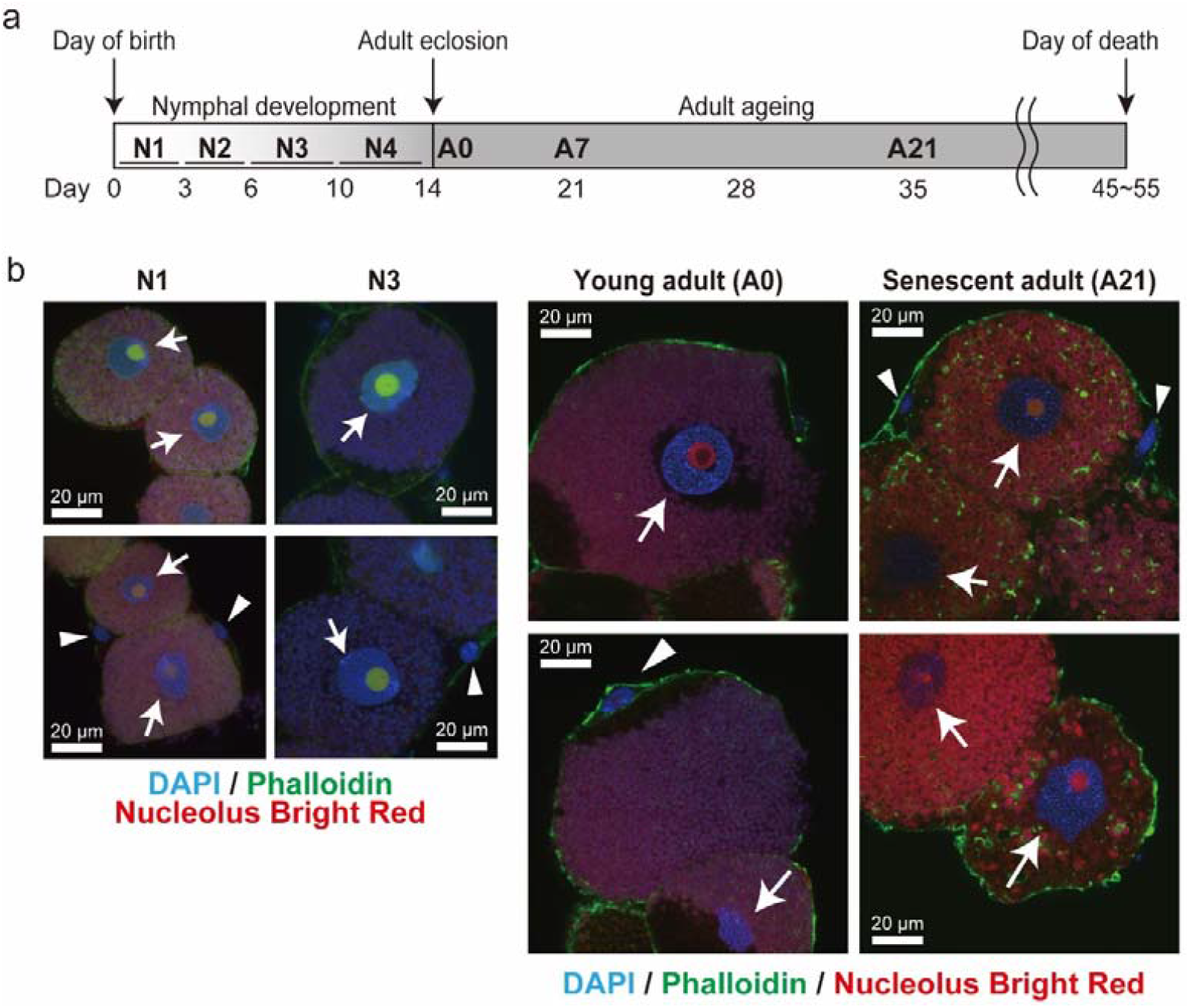
Developmental scheme and morphological changes of bacteriocytes during post-embryonic development in viviparous females. **a** First-instar nymphs of viviparous aphids molt four times during 13.73 ± 0.32 (mean ± SEM, *n* = 16) days at 16 °C. After adult eclosion, aphids live approximately 1 month under 16 °C. N1-4 represents the first to fourth instar nymphs. A0, A7, and A21 mean adult aphids at the day of eclosion, at 7 and 21 days after eclosion, respectively. **b** Confocal microscopic images of bacteriocytes and sheath cells in each stage of viviparous females. N1 and N3 were selected and presented as representatives of nymphal stages. A0 and A21 adults were presented as young adults and senescent adults, respectively. DNA and F-actin were stained using DAPI (blue) and Phalloidin (green), and nucleolus was visualized by Nucleolus Bright Red (red). *White arrows*; nuclei of bacteriocytes, *white arrowheads*; nuclei of sheath cells.

### Ploidy dynamics of aphid bacteriocytes along with post-embryonic development

During post-embryonic development of viviparous females, the ploidy level of bacteriocytes gradually increased; bacteriocytes were 16-32C at the time of birth (N1) and reached the highest ploidy level in actively reproducing adults (A7, 128-256C) (Figure 7). All stages of viviparous females except senescence stage A21 showed significant differences in ploidy levels (Brunner–Munzel test with Bonferroni adjustment, N1 < N2 < N3 < N4 < A0 < A21 < A7, *p* < 0.05, Figure 7). The highest dominant ploidy class was observed in A7 aphids (256C, 43%) (Figure 7). A similar pattern was observed in oviparous aphids (Figure S9), and the highest dominant ploidy was also observed in A7 individuals (but 128C, 47%).

**Figure 7.**
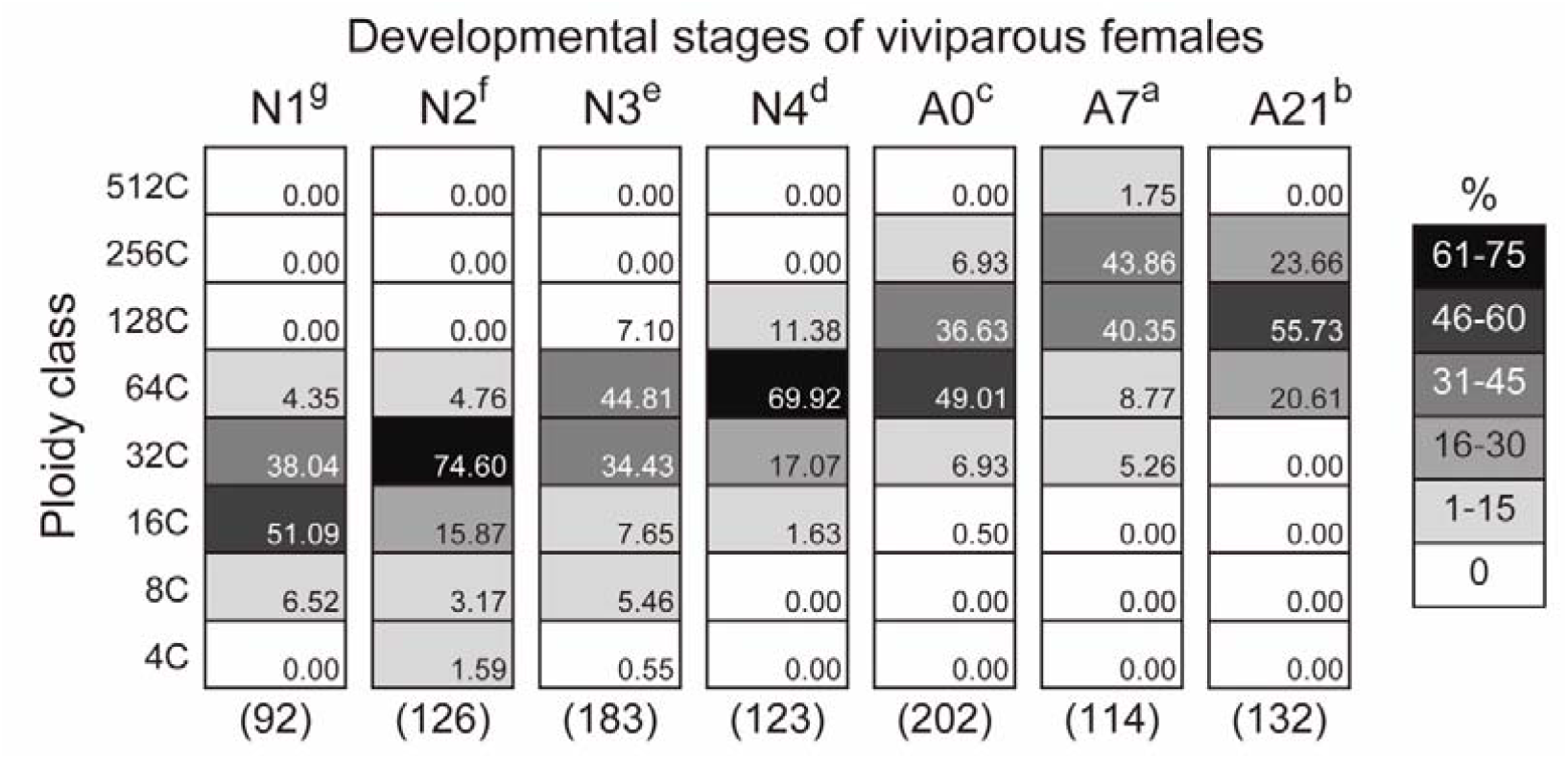
Ploidy distribution of bacteriocytes of each developmental stage of viviparous aphids. Sample size (numbers of bacteriocyte nuclei) is shown below each column. N1-4 represents the first to fourth instar nymphs, and A0, A7 and A21 indicate 0-, 7-, 21-day-old adults. Different letters with aphid stages indicate significant differences in the median ploidy class (Brunner–Munzel test with Bonferroni adjustment, *p* < 0.05). Bacteriocytes of adult viviparous females (A0-A21) in the pea-aphid were highly polyploid (64-256C), while those of N1 nymphs were mainly 16C and 32C cells.

### Relationship between the size of the nucleolus and ploidy levels in aphid bacteriocytes

There were significant effects of ploidy class on the size of the nucleolus in adult bacteriocytes of each morph (LM with type II test; viviparous females, *F* = 62.94, *df* = 2, *p* < 0.001, oviparous females, *F* = 23.97, *df* = 2, *p* < 0.001; males, *F* = 6.44, *df* = 3, *p* < 0.001, Figure 8a). Note that 16C and 256C viviparous bacteriocytes were excluded from the analysis due to their small number. Similarly, 16C and 8C cells of females and males, respectively, were excluded from the analysis. In viviparous females, the size of the nucleolus consistently increased from 32C to 128C (Tukey’s test, *p* < 0.001). In oviparous females, 128C cells had larger nucleoli than 32C and 64C cells (*p* < 0.001 each), yet the difference between 32C and 64C cells was marginally non-significant (*p* = 0.06). In males, the size of the nucleolus of 128C cells was significantly larger than that of 16C and 32C cells (*p* < 0.001 each), but we did not find any significant difference among other comparisons (16C vs. 32C, *p* = 0.71, 16C vs. 64C, *p* = 0.10, 32C vs 64C, *p* = 0.08, 64C vs 128C, *p* = 0.18) (Figure 8a). A significant effect of ploidy class on the nucleolus size was also detected in the data from each developmental stage of viviparous aphids (LMM with type II test; χ^2^ = 788.83, *df* = 5, *p* < 0.001, Figure 8b). Note that 4C and 512C bacteriocytes were excluded from the analysis because of the small number of observations. The size of the nucleolus consistently increased from 8C to 256C (Tukey’s test, *p* < 0.05, Figure 8b).

**Figure 8.**
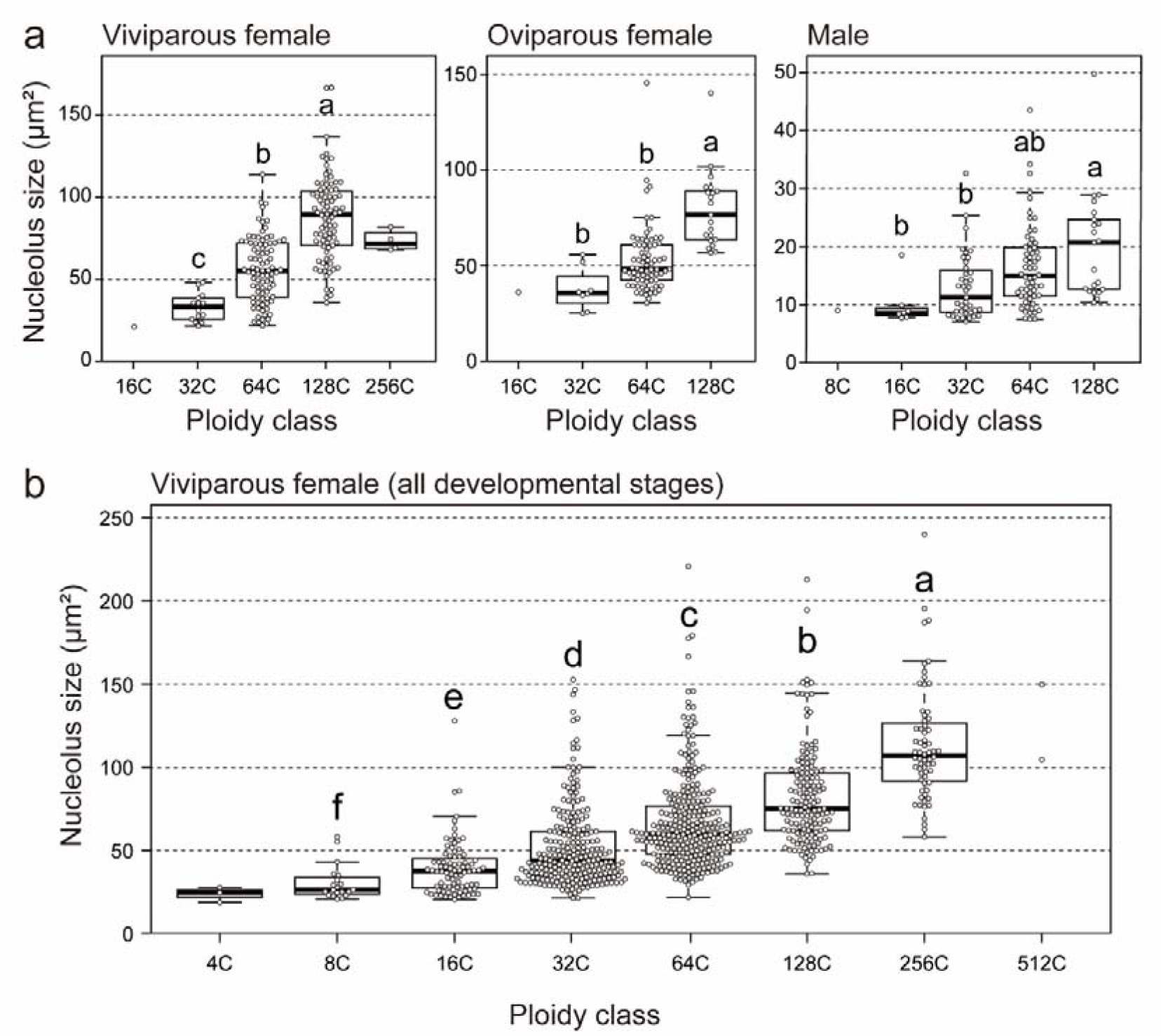
The relationship between the size of nucleolus and ploidy levels of aphid bacteriocytes. In the boxplots, central bold lines represent the medians, boxes comprise the 25–75 percentiles and whiskers denote the range. **a** The size of nucleolus was significantly different among ploidy class in all aphid morphs (LM with type II test; *p* < 0.001 each). Different letters indicate significant differences (Tukey’s test, *p* < 0.05). Note that 16C/256C, 16C, and 8C bacteriocytes in viviparous, oviparous females and males, respectively, were excluded from these analyses due to the small sample size. **b** Nucleolus size was significantly different among ploidy class in viviparous females (LMM with type II test; *p* < 0.001). Different letters indicate significant differences (Tukey’s test, *p* < 0.05). Note that 4C and 512C were excluded from this analysis due to the small number.

## Discussion

This study presented quantitative data on ploidy levels and ploidy dynamics in the bacteriocytes, which are pivotal cells in aphid/*Buchnera* endosymbiosis. The method developed for ploidy analysis of aphid bacteriocytes in this study (Figure 4) revealed the hyper polyploidy of the bacteriocytes ranging from 16- to 256-ploidy throughout the lifecycle. We also found significant differences in the ploidy levels among morphs in adult stages (viviparous females > oviparous females > males, Figure 5) and developmental stages in viviparous females (reproducing adults > senescent adults > pre-reproducing adults > nymphs, Figure 7). Considering that viviparous adults exhibited a high rate of reproduction, which was at a maximum in the first three weeks (Figure S6), more metabolically active (actively reproducing) individuals may show higher polyploidy in their bacteriocytes. We observed a similar pattern of bacteriocyte polyploidy in oviparous aphids (Figure S9), yet the highest ploidy level was lower than viviparous females. These results provide fundamental information to understand the functional significance of polyploidy in aphid bacteriocytes.

Our findings in this study raise the possibility that bacteriocyte polyploidy can enhance nutritional symbiosis between aphids and *B. aphidicola*, wherein both supply each other with nutrients that they cannot synthesize on their own^[10, 19, 20]^. Specifically, we predict that genes involved in amino acid metabolism, related to transport, and for symbiont regulation (e.g., lysozymes or cysteine-rich secreted proteins targeting bacteria), which have been reported to be highly expressed in the aphid bacteriomes^[24, 25, 38, 40]^, may be upregulated in a ploidy-dependent manner^[6, 7, 47]^. On the other hand, it is commonly known that cell volumes change depending on ploidy levels ^[6, 48]^. In fact, our data suggested a positive correlation between bacteriocyte volume and ploidy level in the aphid bacteriocytes (Figure S5 and S7). Bacteriocyte enlargement may simply increase the number of *B. aphidicola* that can be harbored, which should lead to enhanced nutritional symbiosis. In order to examine the effect of polyploidization on the aphid/*Buchnera* symbiosis, an integrated approach including gene expression analysis on a per-nucleus basis, and monitoring cell phenotypes such as ploidy-level and cell/nuclear size is required.

In this study, we also found positive correlations between the ploidy class of bacteriocytes and the size of their nucleoli, regardless of the morphs and developmental stage (Figure 8). The main function of the nucleolus is ribosomal biogenesis, and the size and morphology of nucleoli are linked to nucleolar activity, such as transcription and ribosomal RNA production rates^[49–51]^. In plants, there is evidence that more polyploid nuclei not only exhibit larger nucleolar size but also exhibit increased transcription of rRNA and mRNA; a positive correlation between DNA content and transcriptional activity has been identified in the polyploid tomato fruit pericarp^[8]^. Our data are consistent with the hypothesis that aphid bacteriocytes with higher ploidy levels produce more ribosomal RNA, leading to higher metabolic activity; a direct relationship between ploidy levels and nucleolar activity needs to be examined in future studies.

Aphid bacteriocytes have long been considered to be polyploid [*Myzus persicae* (Sulzer)^[18]^, *Pemphigus spyrothecae* Passerini^[22]^, and *Cinara* species^[52]^]. Furthermore, it has been concluded that bacteriocytes are polyploid in many other insects, such as psyllids, whiteflies, scale insects, weevils, bark beetles, termites, and cockroaches^[11, 12, 15, 53–56]^, although quantitative data were lacking, except for psyllids^[15]^. In this study, by comprehensively describing the patterns of polyploidization, we demonstrated, for the first time, that high metabolic demand such as active reproduction is associated with higher polyploidy levels in aphid bacteriocytes. It would be valuable to investigate this relationship in intracellular symbiosis in various insect species. Accumulating information on bacteriocyte polyploidy will help us gain a better understanding of the maintenance and evolution of mutual relationships between host and symbionts because polyploidization in the symbiotic host cells is a common rule in insect-microorganism intracellular symbioses^[12, 13, 57]^.

In the present study, we compared three methods for ploidy level quantification and found an image-based fluorometry the best for the analysis of aphid bacteriomes (Figure 4), because it could distinguish between cell types (e.g., bacteriocytes and sheath cells) by the size of nucleolus, unlike flow cytometry (Figure S2). In addition, it was timesaving compared with Feulgen densitometry (Figure S3). Our results from fluorometry showed that sheath cells mainly consisted of 16C and 32C in adult viviparous aphids, although peaks in the histogram were not clear (Figure 4); therefore, nuclei exhibiting more than 32C can be reasonably assumed to be bacteriocytes. These approach, the combination of several methods such as the fluorometry and flow cytometry would be applicable to other symbiotic systems of insects with microorganisms, wherein the bacteriome frequently contains several types of cells (e.g., primary or secondary bacteriocytes, and sheath cells)^[13, 14, 21]^.

In conclusion, we comprehensively described the patterns of polyploidization in aphid bacteriocytes, which has long been assumed to be polyploid, yet there have been no quantitative studies ^[9, 10, 14]^. Based on the patterns and cytological features observed in this study, we suggest that hyper polyploidy may enhance gene expression levels and increase cell size, contributing to the nutritional symbiosis with the bacterial symbiont *B. aphidicola*. This study provides a foundation for further molecular-level analysis of the functions and underlying mechanisms of polyploidy in insect symbiotic host cells.

## Material and methods

### Aphids

In this study, we used a long-established parthenogenetic clone of the pea-aphid, *A. pisum*, ApL strain, which was originally collected in Sapporo, Hokkaido, Japan (referred to as Sap05Ms2 in ^[58]^). We confirmed that this strain only harbors the primary endosymbiont *B. aphidicola* by diagnostic PCR, as described in a previous study^[59]^. Viviparous insects were maintained on young, broad bean plants (*Vicia faba* L.) in a 16 °C incubator, with a photoperiod of 16 h light:8 h dark (long-day conditions). Sexual morphs (oviparous females and males) were induced by short-day conditions (e.g., 8 h light:16 h dark) (Figure 2, S1, modified from ^[58, 60, 61]^). Viviparous females were randomly selected from the synchronized source populations and reared on young broad bean plants at 16 °C under long-day conditions. Newly larviposited aphids (G0, first-instar nymphs, Figure S1) were then transferred onto the leaves of the bean plants in a 15 °C incubator with a photoperiod of 8 h light:16 h dark. After these G0 nymphs grew into adults, they started to produce G1 offspring, which are morphologically identical to apterous/viviparous females but produce sexual individuals (Figure S1). G1 adults produce G2 offspring that contains sexual individuals (oviparous females and males), but they also produce a few viviparous females (Figure S1). In this study, all oviparous females and males were G2 offspring, and all viviparous individuals were apterous. To reveal the fecundity and longevity of both viviparous and oviparous aphids, female aphids were reared separately and observed daily (see supplementary information “SI Methods”).

### Size and morphology of aphid bacteriome cells

Aphid bacteriomes consist of two types of cells: bacteriocytes containing *Buchnera* cells in their cytoplasm and sheath cells without them (Figure 1b, ^[9, 21, 40]^). To gain more detailed cellular features of aphid bacteriocytes, we performed a morphological analysis of bacteriomes using a confocal laser-scanning microscope (CLSM; FV1000, Olympus, Japan) on three adult morphs (viviparous/oviparous females and males). These adults were within 5 days of eclosion. We also observed the bacteriome at each stage of viviparous females [nymphs N1 (first-instar nymph), N2 (second-instar nymph), N3 (third-instar nymph), and N4 (fourth-instar nymph), and young adults (3-5 days after eclosion), and old adults (approximately 21 days after eclosion)]. Each stage of viviparous females was used on the day of molting, but teneral insects were not used). Aphid bacteriomes were dissected in phosphate-buffered saline (PBS: 33 mM 143 KH_2_PO_4_, 33 mM Na_2_HPO_4_, pH 6.8) under a stereomicroscope (SZ61, Olympus, Japan), with fine forceps, and their bacteriome cells (bacteriocytes and sheath cells) were surgically isolated from the aphid abdomen. The cells were fixed with 4% paraformaldehyde in PBS for 30 min. Fixed bacteriome cells were washed three times in 0.3% Triton X-100 in PBS (PBS-T) for 15 min for permeabilization. The cells were then stained with 4,6-diamidino-2-phenylindole (DAPI) (1 μg/mL; Dojindo, Japan) for the nuclei and Alexa Fluor™ 488 phalloidin (66 nM; Thermo Fisher Scientific, USA) for the cytoskeleton (F-actin), respectively. The nucleolus, which is the site of both ribosomal RNA (rRNA) synthesis and the assembly of ribosomal subunits^[49]^, was visualized using Nucleolus Bright Red (1 mM; Dojindo, Japan). Nucleolus Bright Red dyes are small molecules that electrostatically bind to RNA in the nucleolus to emit fluorescence. After 1 h at room temperature (from 20 to 25 °C), the cells were washed three times with PBS-T for 15 min and mounted with VECTASHIELD antifade mounting medium (Vector Laboratories, USA). The morphology of the bacteriocytes and sheath cells was visualized using fluorescent staining and differential interference contrast microscopy. The captured images were processed using the image analysis software ImageJ (NIH, http://rsb.info.nih.gov/ij/). The diameter of each bacteriocyte (D) was measured at its widest point. The approximate bacteriocyte volume (V) was calculated using the standard formula: 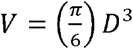. The nucleolus size (area) of both the bacteriocytes and sheath cells was also recorded.

### Establishment of the method for ploidy analysis on aphid bacteriocytes

To establish a method for ploidy analysis of aphid bacteriocytes, we used three methods: flow cytometry, Feulgen densitometry, and fluorometry. Young adult viviparous aphids (3-5 days after eclosion) or late instar (third-fourth instar) of nymphs were used. In all analyses, not only bacteriome cells but also head cells were used as diploid controls, which were confirmed as diploid by preliminary analysis of sperm cells (haploid). We first performed flow cytometry, which is an efficient and commonly used method for nuclear DNA-content analysis^[62, 63]^. For this analysis, bacteriome cells dissected from five aphids were suspended and repeatedly pipetted in 250 μL of trypsin buffer (0.11% Nonidet P40, 0.1% sodium citrate, 0.05% spermine tetrahydrochloride, 0.01% Tris base, and 0.003% trypsin in distilled water). Heads were ground with tight-fitting pestles in the trypsin buffer. After incubation at room temperature (from 20 to 25 °C) for 10 min, 5 μL of trypsin inhibitor solution (trypsin inhibitor, from soybean, 25 mg/mL) and 1 μL of RNase A (100 mg/mL) were added. After the mixture was incubated at room temperature (from 20 to 25 °C) for 10 min, 250 μL of dilution buffer (0.11% Nonidet P40, 0.1% sodium citrate, 0.17% spermine tetrahydrochloride, and 0.01% Tris base in distilled water) was added, and the mixture was filtered through a 48 μm nylon mesh. Isolated nuclei were stained with DAPI (1 μg/mL). The filtered mixture was incubated at room temperature for at least 10 min and stored on ice until use. Stained nuclei were analyzed for DNA-DAPI fluorescence using a Cell Sorter SH800 (SONY, Japan) at an excitation wavelength of 405 nm. Each experiment was performed in triplicates.

Second, we conducted Feulgen densitometry, which have been widely used for DNA content analysis ^[64]^. This method was used in the ploidy analysis of psyllid bacteriome cells ^[15]^. This analysis was performed according to the protocol of Hardie *et al*. (2002)^[64]^. Briefly, the bacteriome cells and heads were dissected from an aphid and then smeared on glass slides. The smears were fixed in MFA (methanol, formalin, acetic acid = 85:10:5 v/v) for 24 h, hydrolyzed in 5.0 N HCl for 2 h, and stained with Schiff reagent for 2 h. Images of stained nuclei were captured with a BX-61 microscope (Olympus, Japan) and a DS-Fi1 CCD camera (Nikon, Japan). All the steps were performed at room temperature (from 20 to 25 °C). Using ImageJ, the green channel was extracted, and the integrated optical density (IOD) of the Feulgen stain in the nuclei was measured. Background signal intensity was measured in an area adjacent to each nucleus and deduced from the nuclear IOD. Each experiment was performed in triplicates. In this analysis, the nuclei of bacteriome cells were not isolated; therefore, we can categorize cell types based on their cytoplasmic status [*Buchnera* presence (bacteriocyte) or absence (sheath cells)].

Third, image-based fluorometry of the isolated nuclei was conducted. Bacteriome cells were dissected from three individuals in PBS, and the PBS droplets containing the cells were transferred onto glass slides by careful pipetting. Nuclear isolation buffer (10 μL), identical to “dilution buffer,” was added to the cells on the glass slide. The nuclei of these cells were isolated by agitation using fine needles (insect pins, Shiga Konchu Fukyusha, Japan) in the droplets. This step was completed using an SZ61 stereomicroscope. After drying the nuclei on the glass slide, they were fixed in MFA for 30 min at room temperature (from 20 to 25 °C). The slides were washed thrice with distilled water. Nuclei were stained with DAPI solution (1 μg/mL DAPI and 2 mg/mL RNase A) for 1 h at room temperature (from 20 to 25 °C). The slides were washed three times with distilled water and mounted with VECTASHIELD antifade mounting medium. The same devices used for the Feulgen densitometry were used for image capturing. Using ImageJ, the blue channel was extracted, and the integrated fluorescent intensity (IFI), which is the integrated gray value in the region of interest, was measured for each nucleus. The background signal intensity was measured in an area adjacent to each nucleus and deduced from nuclear IFI. The experiment was performed in duplicates. In this analysis, the nucleolus size (area) was also measured and compared with the data from confocal microscopy to discriminate cell types (bacteriocytes and sheath cells).

### Ploidy analysis for each morph, and each developmental stage of viviparous aphids

Bacteriome cells and heads were dissected in PBS and processed with the abovementioned “fluorometry” method. First, to assess the variation in the degree of polyploidy among morphs, young adult individuals (3-5 days after adult eclosion) of viviparous/oviparous females and males were used. Three individuals were pooled for each morph. Nucleolus sizes were recorded to discriminate cell types. Second, to elucidate the dynamics of polyploidization and post-embryonic development of aphids, bacteriocytes of viviparous aphids at different developmental stages were analyzed. Specifically, the following stages were used: nymphs N1, N2, N3, and N4, and adults at three distinct time points: A0 (0 days after eclosion) as pre-reproductive adults, A7 (7 days after eclosion) as actively reproducing adults, and A21 (21 days after eclosion) as senescent individuals. All viviparous individuals were apterous. Each stage of viviparous females was used on the day of molting, but teneral insects were not used. To discriminate cell types, the nucleolus size (area) was measured and compared with the data obtained by confocal microscopy. Nucleolus sizes have also been used as indicators of the translation activity of cells ^[8, 49]^. Three individuals were used in each stage. Additionally, each developmental stage of oviparous females (nymphs N1, N2, N3, and N4, and A0 and A7 adults) was also analyzed. For discrimination of cell types in oviparous females, data from viviparous females were used, because we preliminarily confirmed that the size of the nucleolus of bacteriocytes in oviparous females was not significantly different from that in viviparous females. Three individuals were included in each stage.

### Statistical analysis

To compare the size of bacteriocytes among aphid morphs (young adults of viviparous females, oviparous females, and males), we used a linear model (LM), followed by Tukey’s post hoc test. In this analysis, morphs were treated as fixed effects. For analysis of cell size among the developmental stages of viviparous females, we used linear mixed models (LMM), followed by Tukey’s post hoc test. In the analysis, developmental stages and individuals were included as fixed and random effects, respectively. For pairwise comparisons of ploidy levels of bacteriocytes among morphs and developmental stages of viviparous/oviparous females, we used the Brunner–Munzel test, which is a non-parametric test that adjusts for unequal variances. Significant *p-*values were adjusted using Bonferroni’s correction. To compare the size of the nucleolus in bacteriocytes, which was recorded during fluorometry, among ploidy classes, we used LMs and LMMs, followed by Tukey’s HSD post hoc test. In these analyses, ploidy classes were treated as factorial fixed effects. All analyses were performed for each morph. In the analysis of developmental stages, data from each stage were pooled, and individuals were included as a random effect. All analyses were conducted using the “car”, “emmeans”, “lawstat”, “lme4”, and “multcomp” packages in R software v4.1.1 (https://www.r-project.org/).

## Acknowledgements

We thank S. Yorimoto, C. Chung, M. Suzuki, and other members of the laboratory of Evolutionary Genomics, NIBB for their assistance and valuable discussions. We also thank A. Nakabachi and S. Koshikawa for technical advice, Functional Genomics Facility, NIBB Core Research Facilities for the technical support, and Editage (www.editage.com) for English language editing. This work was financially supported by the Japan Society for the Promotion of Science to T.N. (Research Fellowship for Young Scientists No. 19J01756) and S.S. (KAKENHI 17H03717 and 20H00478).

## Author contributions

T. N. and S. S. designed research; T. N. performed the experiment and analyzed data; T. N. and S. S. wrote the original draft of the paper and both authors will contribute substantially to revisions.

