## Supplemental Info for "Ploidy dynamics in aphid host cells harboring bacterial symbionts"

Original Research Article

Supplementary Information

Tomonari NOZAKI^1*^, Shuji SHIGENOBU^1,2^

*^1^Laboratory of Evolutionary Genomics, National Institute for Basic Biology, 38 Nishigonaka, Myodaiji, Okazaki, Aichi 444-8585, Japan*

*^2^Department of Basic Biology, School of Life Science, The Graduate University for Advanced Studies (SOKENDAI), 38 Nishigonaka, Myodaiji, Okazaki, Aichi 444-8585 Japan*

^*^*Correspondence*:

Tomonari Nozaki, Laboratory of Evolutionary Genomics, National Institute for Basic Biology, Okazaki, Aichi 444‐8585, Japan


SI Methods

*Productivity and developmental schemes of female aphids*

To compare the productivity and longevity between viviparous and oviparous aphids, newly molted adults were chosen and separately reared on the leaves of the bean plants, *Vicia faba*. Both females were apterous. Oviparous females were reared with young males from the same generation, allowing for free mating. These females were checked daily, and both the number of larviposition (the number of nymphs produced) by viviparous females and oviposition (the number of eggs produced) by oviparous females were recorded. The day of death was also recorded. Viviparous adults were maintained at 16 °C under a long-day photoperiod (16 h light:8 h dark), the same condition as the source populations. Oviparous adults were placed at 15 °C under a short-day condition (8 h light:16 h dark). All nymphs and eggs laid were removed immediately after the daily observation to avoid crowded conditions, which can influence female development ^[1]^ and induce oocyte/embryo resorption ^[2]^. To elucidate the developmental scheme (duration of each stage at 16 °C) of viviparous females, newly larviposited (first instar) nymphs were reared separately and checked daily to record the day of molting.

We compared the total number of larvae and eggs laid during the entire life of insects using a generalized linear mixed-effect model (GLMM) with a Poisson error distribution and log link function between viviparous and oviparous females. In this analysis, the types of females were treated as fixed effects, and individuals were included as a random effect. Longevity (= adult lifetime) of both types of females was also compared. In this model, the types of females were treated as a fixed effect, and individuals were included in a random effect. Note that the interaction between female types and their lifespan was analyzed but not significant (see Results).

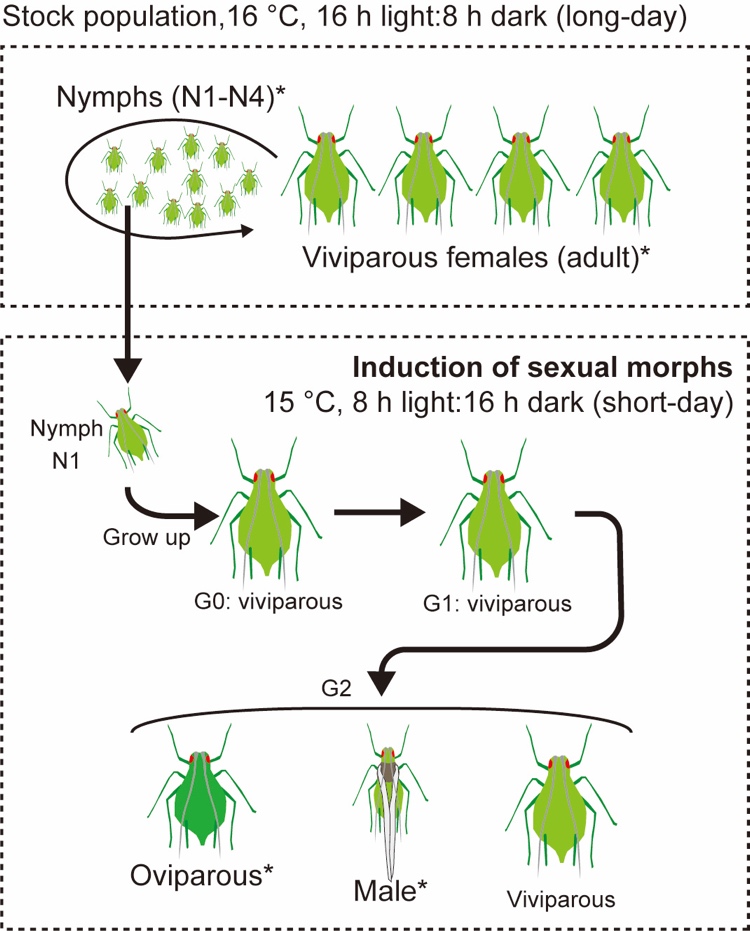
Supplementary Figures

**Figure S1.** Detailed experimental procedure to induce males and oviparous females in the ApL strain. In the stock population, viviparous insects were maintained on young, broad bean plants (*Vicia faba* L.) under 16 °C, a photoperiod of 16 h light:8 h dark (long-day conditions). First instar nymphs isolated from stock populations (G0 insects) were reared on the broad bean plants, under 15 °C, 8 h light:16 h dark (short-day condition). Offspring of them (G1 insects) parthenogenetically produce sexual morphs (G2, Oviparous females and males). G1 aphids also produce few viviparous females in the ApL strain.

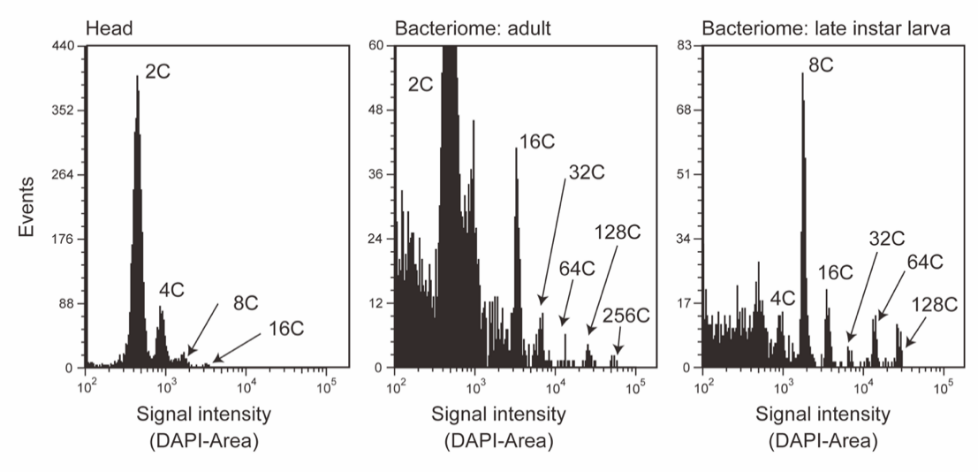

**Figure S2**. Examples of nuclear DNA content analysis by flow cytometry. The first peak in the head sample corresponds to the population of 2C-DNA nuclei, which was preliminarily determined by analysis of sperm cells (haploid; 1C) of males (data not shown). Analysis on bacteriome of adults and late instar nymphs revealed that this organ consisted of at most 256C cells, while cell types could not be identified. In adult bacteriome analysis, a large number of 2C nuclei were occasionally observed, probably due to the contamination of embryonic cells.

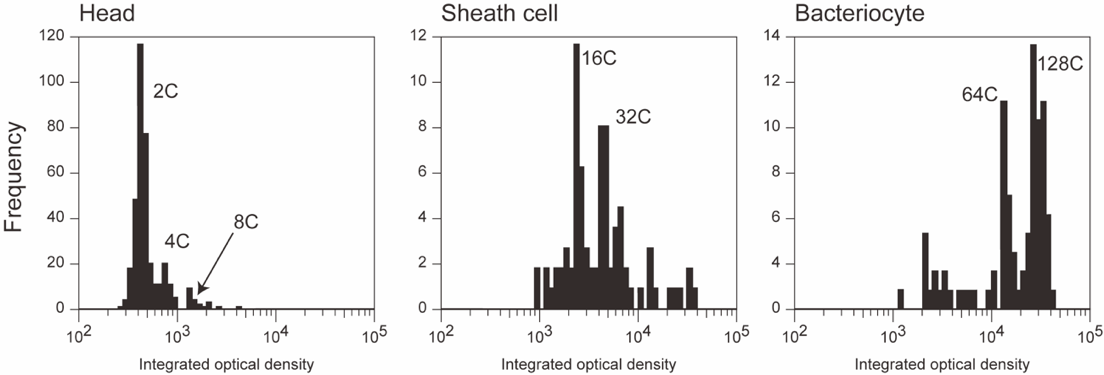

**Figure S3**. Examples of nuclear DNA content analysis by Feulgen densitometry. The first peak in the head sample corresponds to the population of 2C-DNA nuclei, which was preliminarily determined by analysis of sperm cells (haploid; 1C) of males (data not shown). Analysis of the bacteriome cells of adult viviparous aphids demonstrated that these bacteriocytes and sheath cells mainly consist of 64C and 128C, and 16C and 32C cells, respectively, although the cell numbers were small and each peak was not clearly separated.

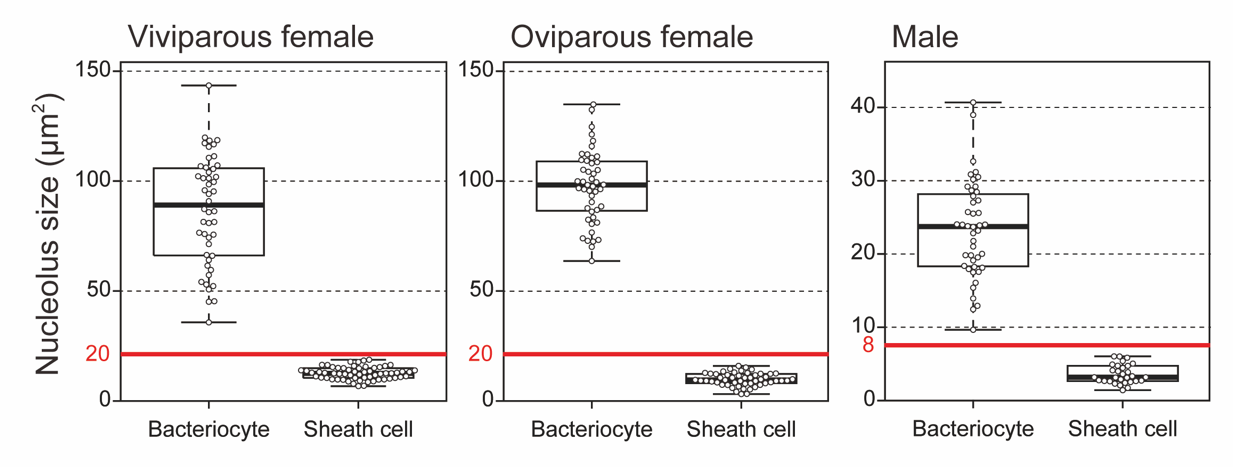

**Figure S4.** Nucleolus size distribution in three types of adults: viviparous females, oviparous females, and males. The size of nucleolus was significantly different between bacteriocytes and sheath cells, regardless of aphid morphs (LM with type II test, *p* < 0.001 in all). There was no overlap in nucleolus sizes between the two types of cells. Red bars indicate the “threshold” used in the image-based fluorometry (see Material and methods).

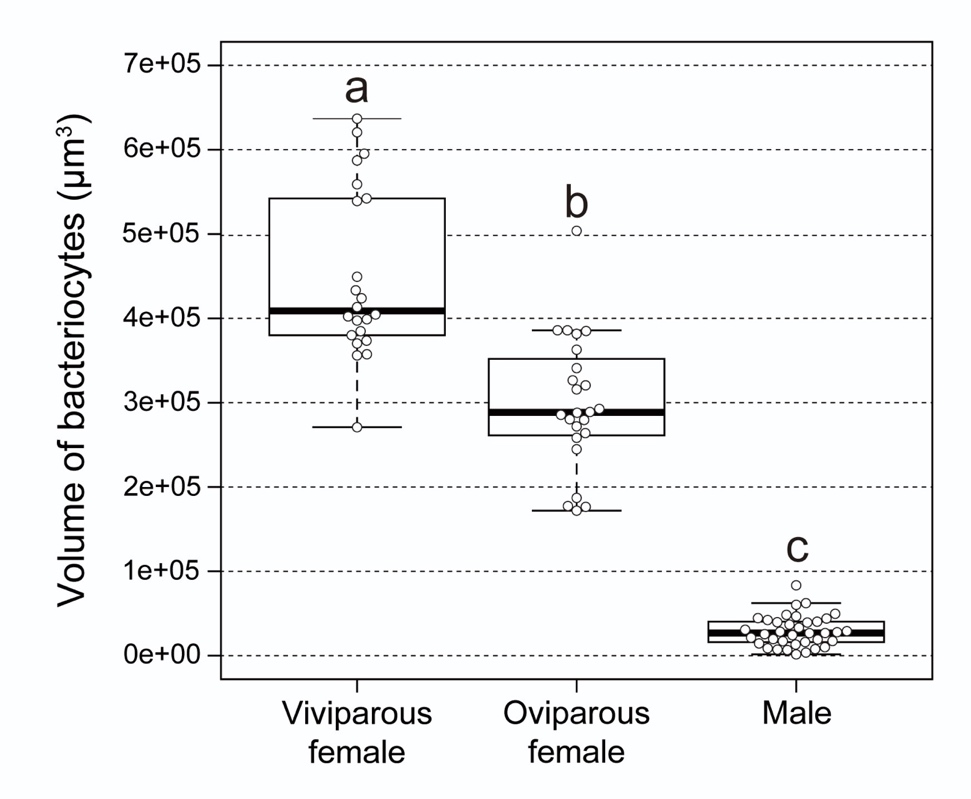

**Figure S5.** Cell volume distribution of bacteriocytes in each aphid morph. The volume of bacteriocytes was estimated based on the data from confocal microscopy (see Material and methods). Results are displayed as boxplots where central bold lines represent the medians, boxes comprise the 25–75 percentiles, and whiskers denote the range. The size of bacteriocytes was significantly different among aphid morphs [viviparous females; 449847.05 ± 21583.0 (mean ± SEM) μm^3^, *n* = 22, oviparous females; 298989.9 ± 16196.6 μm^3^, *n* = 24, and males; 29020.35 ± 3001.6 μm^3^, *n* = 37] (LM with type II test, *p* < 0.001). Different letters indicate significant differences (Tukey’s test, *p* < 0.05).

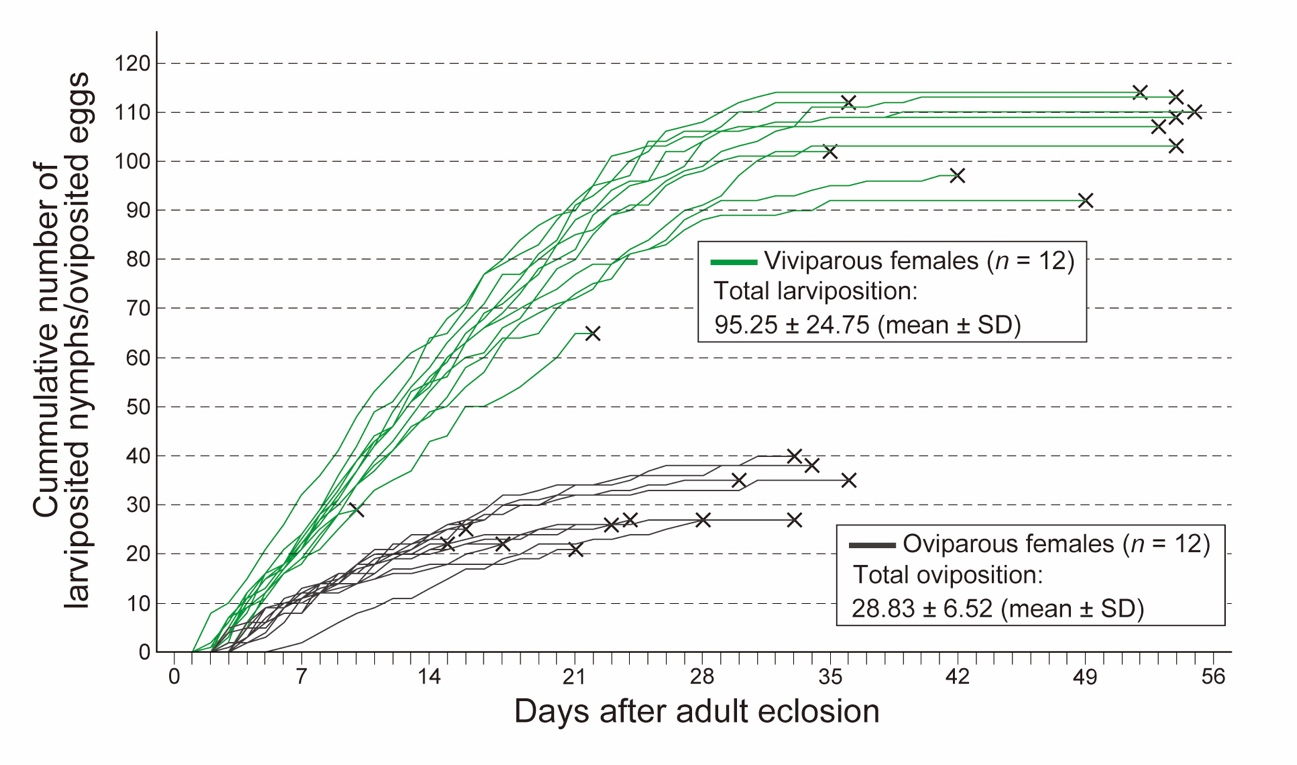

**Figure S6.** Reproduction activity in both types of aphid females. Viviparous aphids start larviposition several days after adult eclosion and reproduce actively during approximately 3 weeks. In total, the number of larviposition was approximately 90-100 in viviparous females [95.25 ± 24.75 (mean ± SD)], while lifetime oviposition by oviparous females was less than one-third of larviposition by viviparous females [28.83 ± 6.52 (mean ± SD)]. Cross marks indicate the death of the insect. Viviparous females and oviparous females were maintained on the leaves of bean plants, *Vicia faba*, under 16 °C, 16 h light (L):8 h dark (D) and 15 °C, 8L:16D, respectively. Oviparous females were maintained with adult males (these males were removed after their death). Viviparous females started reproduction from days 2-3 and the rate of larviposition was at the peak during days 3-20 but slow downed during days 21-28. They lived at most 50-55 days, although most of them stopped larviposition after day 30. In oviparous females, first oviposition and mating with males were observed at days 3-4. They laid eggs actively until around day 14 but their death was observed almost simultaneously.

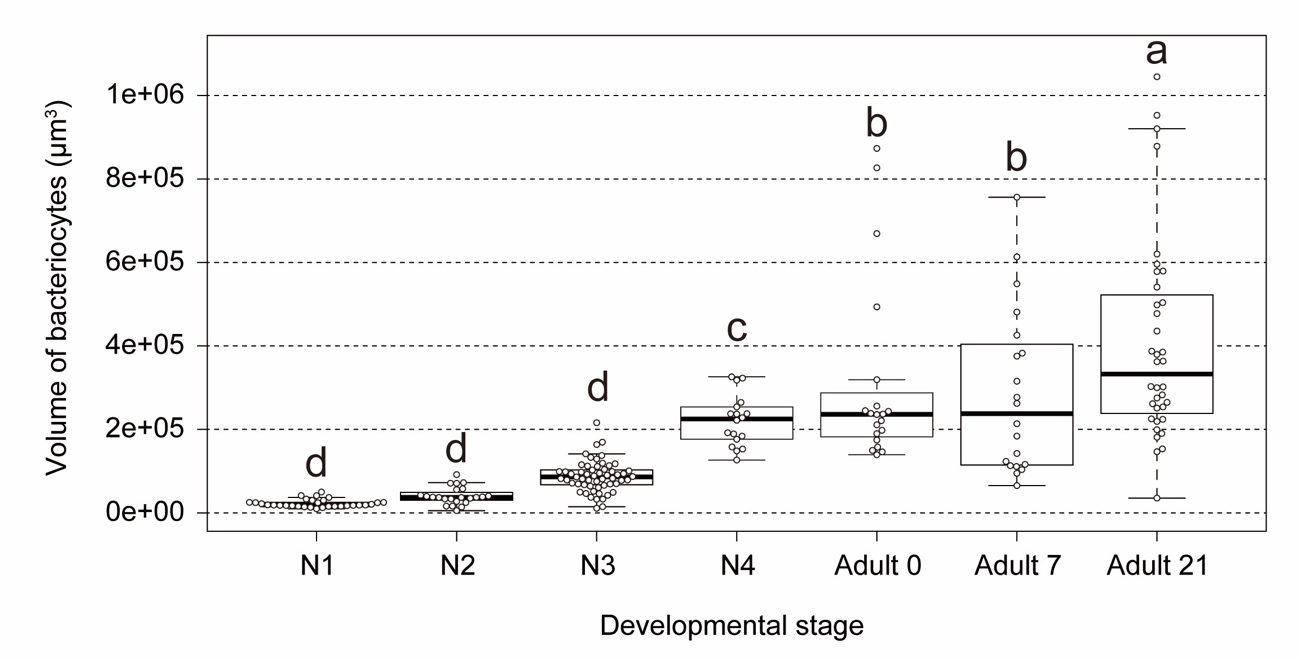

**Figure S7.** Cell volume distribution of bacteriocytes in each developmental stage of viviparous aphids. The volume of bacteriocytes was estimated based on the data from confocal microscopy (see Material and methods). Results are displayed as boxplots, where central bold lines represent the medians, boxes comprise the 25–75 percentiles, and whiskers denote the range. N1-4 represents the first to fourth instar nymphs. The size of bacteriocytes was significantly different among developmental stages [N1; 22765.0 ± 1703.7 (mean ± SEM) μm^3^, *n* = 31, N2; 40665.0 ± 4386.3 μm^3^, *n* = 23, N3; 87142.7 ± 4922.5 μm^3^, *n* = 57, N4; 220822.5 ± 14305.6 μm^3^, *n* = 18, Adult 0; 311178.5 ± 49807.9 μm^3^, *n* = 20, Adult 7; 285287.2 ± 49807.9 μm^3^, *n* = 20, and Adult 21; 404810.9 ± 40292.5 μm^3^, *n* = 36] (LMM with type II test, *p* < 0.001). Different letters indicate significant differences (Tukey’s test, *p* < 0.05).

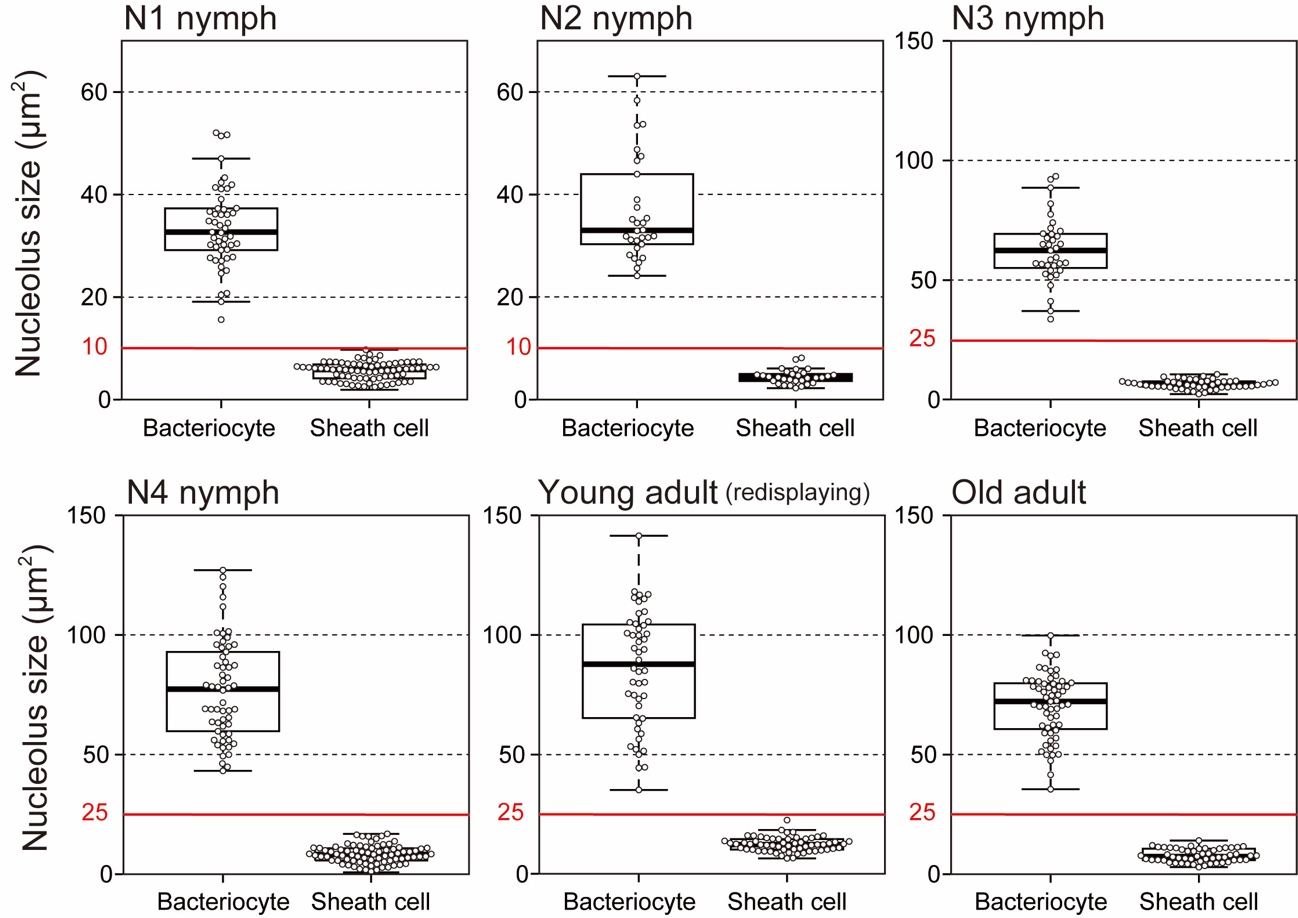

**Figure S8.** Nucleolus size distribution in each developmental stage of viviparous aphids. The size of nucleolus was significantly different between bacteriocytes and sheath cells, regardless of developmental stages (LMM with type II test, *p* < 0.001 in all). There was no overlap in nucleolus sizes between the two types of cells. N1-4 represents the first to fourth instar nymphs. Red bars indicate the “threshold” used in our image-based fluorometry (see Material and methods). Data in the category “Young adults” was the same as “Viviparous females” in Figure S3 (re-displaying).

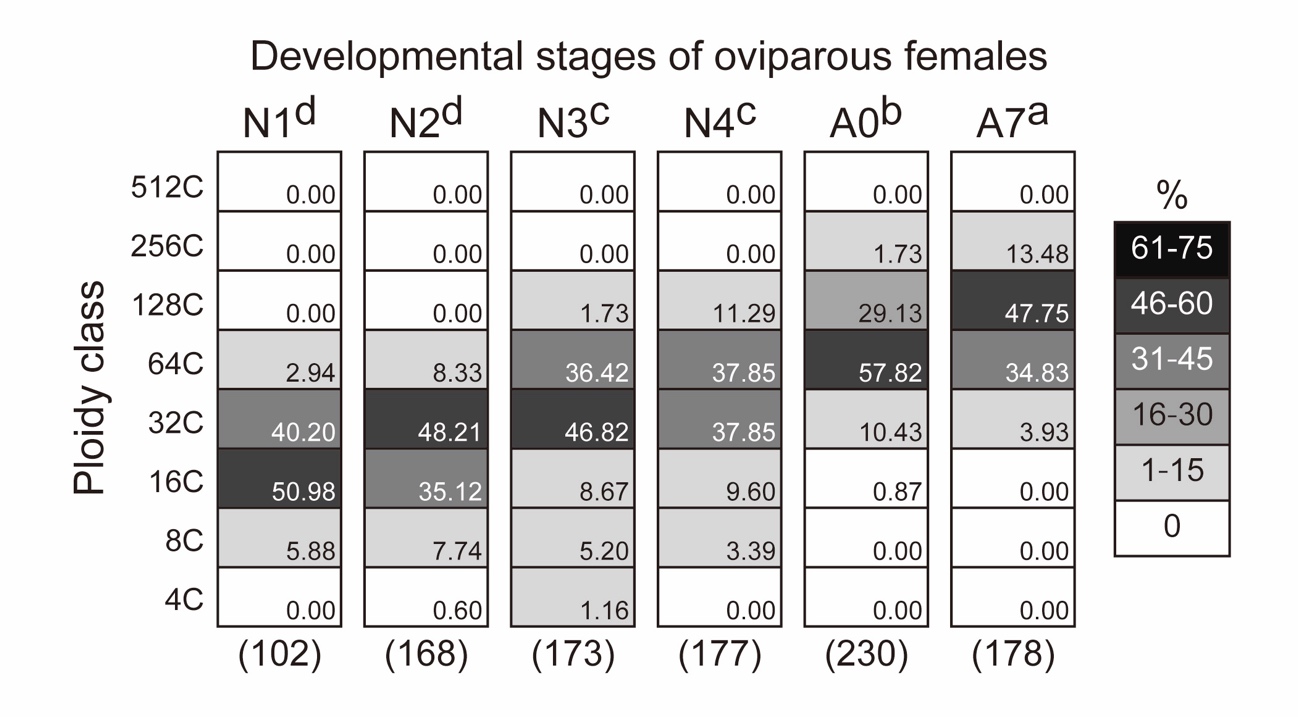

**Figure S9.** Ploidy distribution of bacteriocytes of each developmental stage of oviparous aphids. Sample size (numbers of bacteriocyte nuclei) is shown under the column. N1-4 represents the first to fourth instar nymphs. A0, A7 and A21 indicate 0-, 7-, 21-day-old adults. Different letters with aphid stages indicate significant differences in the median ploidy class (Brunner–Munzel test with Bonferroni adjustment, *p* < 0.05). Bacteriocytes of 7-day-old adults exhibited the highest polyploid level in oviparous aphids (64-128C).

**References (only referred to in the SI)**

1. Sutherland, O. R. W. (1969). The role of crowding in the production of winged forms by two strains of the pea aphid, *Acyrthosiphon pisum*. Journal of Insect Physiology, 15(8), 1385–1410. https://doi.org/10.1016/0022-1910(69)90199-1
2. Ward, S. A., & Dixon, A. F. G. (1982). Selective resorption of aphid embryos and habitat changes relative to life-span. The Journal of Animal Ecology, 859–864. https://doi.org/10.2307/4010
